# Forest Owlet or Farmer’s owlet: Scale-dependent habitat selection reveals conditional compatibility between Forest Owlet conservation and traditional agroforestry in Gujarat, India

**DOI:** 10.64898/2026.02.03.703545

**Authors:** Jenis R. Patel, Kuldeep Gamit, Shashank Patel, Kulbhushansingh Suryawanshi, Anirudhkumar Vasava

## Abstract

Understanding how species that are threatened with extinction utilise human-modified landscapes is essential for evidence-based conservation. We investigated multi-scale habitat selection by the Forest Owlet (*Athene blewitti*), an Endangered species, endemic to central India with fewer than 1000 mature individuals, in the Dangs district of Gujarat, the westernmost extent of its range. Using a hierarchical Bayesian occupancy framework, we examined how forest cover and three agricultural land-use types (dry agriculture with trees, dry agriculture without trees, and intensive agriculture) affected occupancy across three nested spatial scales: regional (81 km^2^), landscape (4 km^2^), and territory (0.25 km^2^). At the regional scale, the forest × agriculture interaction term was significantly negative (β = -6.82, 95% CI: -9.87 to -1.59), indicating that owlets favour agroforestry-dominated regions over forest-dominated landscapes. Conversely, at the landscape scale, a significant positive interaction (β = 1.36, 95% CI: 0.41–2.50) revealed synergistic benefits from forest-agriculture mosaics. Agriculture type strongly influenced landscape-scale occupancy: dry agriculture with trees showed positive effects (β = 1.17, 95% CI: 0.43–2.02), whereas dry agriculture without trees had significant negative effects (β = -1.19, 95% CI: -2.28 to -0.29). These findings demonstrate that Forest Owlets are not forest-obligate specialists but occupy complex agroforestry mosaics, requiring multi-scale conservation strategies. We propose that the traditional Malki agroforestry system, which incentives tree retention on farmland, offers conditional compatibility with Forest Owlet conservation, provided that mature cavity-bearing trees and small forest patches are explicitly protected.

## Introduction

Effective in-situ conservation of rare and endemic species requires a detailed understanding of their habitat requirements at multiple spatial scales (Martínez & Zuberogoitia, 2004; Lipsey et al., 2017). The habitat provides essential resources for survival and reproduction, including refuges, foraging grounds, breeding sites, and corridors for dispersal. Changes in habitat can disrupt these processes. As a result, populations may decline and local extinction can occur. Notably, different life-history processes operate at different spatial scales: foraging takes place within small patches, whereas natal dispersal and meta-population dynamics occur across landscapes. Not considering scale in species habitat relationships can lead to missing key ecological patterns. These patterns are important for population viability and conservation (Šálek et al., 2016; Comfort et al., 2016; Lipsey et al., 2017).

Traditional habitat studies often neglect two critical dimensions. First, landscape configuration matters: the size, connectivity, and spatial arrangement of habitat patches influence species occurrence beyond simple presence or absence of particular land-cover types. Second, habitat effects are inherently compositional, species require multiple landscape elements that together create suitable conditions. A site may provide foraging opportunities but lack nesting substrate, or vice versa. Understanding how different habitat components interact across scales is essential for identifying truly suitable habitat, yet most studies examine variables in isolation at single scales (Lipsey et al., 2017). This approach can misinform management priorities and lead to ineffective conservation designations (Denac et al., 2019).

Agricultural landscapes present particular challenges for habitat assessment because agriculture includes a wide range of practices with very different ecological outcomes. In India, the varied geography means that agricultural systems vary greatly from one region to another, shaped by the climate, topography, history, and policy. Traditional low-intensity farming can provide a habitat for species that depend on forests, whereas mechanised intensive agriculture usually does not. Treating all agricultural land as the same ignores the important ecological differences that may be crucial for conservation planning (Yashmita-Ulman et al., 2018; Aditya & Shaanker, 2020). The Forest Owlet (*Athene blewitti*) is an endemic and endangered owlet that was rediscovered in 1997 after being presumed extinct for over a century (King & Rasmussen, 1998; Rasmussen & Collar, 1998). This owlet is distinct from other Athene species in terms of its morphology and behaviour, with strongly diurnal habits, a direct flight pattern, and a preference for semi-open forest habitats (Ishtiaq & Rahmani, 2005). The current global population is estimated to be between 250 and 999 individuals, which are distributed across fragmented forest patches in central India (BirdLife International, 2018). The species has a consistent association with dry deciduous, teak-bearing forests in the Satpura and adjoining hill systems, showing a marked preference for semi-open, structurally heterogeneous habitats that feature large mature trees with open ground (Kulkarni & Mehta, 2020; Khan et al., 2023). The Forest Owlets nest and roost in the cavities of large, mature trees, which makes the retention of old-growth elements critical for the species’ population to persist (Ishtiaq & Rahmani, 2005; Jathar & Rahmani, 2012).

Recent surveys have revealed an intriguing paradox: despite being classed as a forest-dependent species, Forest Owlets are most often found in agricultural landscapes rather than continuous forest. Previous studies have recorded 82 individuals in south Gujarat, of which 71.4% were in teak-dominated agricultural landscapes, 20.4% in degraded forest, 6.1% in riverine habitat, and only 2.0% in intact forest (Patel et al., 2015, 2017). Also, subsequent study found that the species’ occupancy was positively influenced by both the density of large trees and the cover of agricultural land, with the species occurring where these habitats form a mosaic (Khan et al., 2023). Studies from other regions have found similarly that the species prefers semi-open teak dry deciduous forest interspersed with agricultural fields, while also confirming that it is not found in non-forest habitats, such as pure agriculture and human habitation (Kulkarni & Mehta, 2020). These studies have established Forest Owlets as edge specialists that utilise the interface between forest and agriculture, yet a fundamental question remains unanswered: do Forest Owlets respond to the heterogeneity of agricultural landscapes, or does the forest context modify the effects of agriculture at different spatial scales?

The Dangs District of Gujarat is a prime example of agricultural heterogeneity within a forest matrix. The ‘Malki’ agroforestry system is unique to this area: government policy encourages farmers to grow and protect valuable trees, mainly teak and Terminalia species, along field boundaries, and in return, they are granted ownership rights. These trees cannot be cut down without permission from the Forest Department, which leads to the creation of persistent agroforestry mosaics (Dasa et al., 2022). The hilly terrain and rocky soils in the region cause water scarcity during certain times of the year, despite the area receiving the highest rainfall in Gujarat, resulting in most areas only being able to support one crop a year and fields being left fallow for long periods potentially creating a habitat for Forest Owlets to forage. This mix of forest and agriculture, with large forest areas among villages and farmland, is a rare landscape type in the species’ range. The recent designation of The Dangs as an Aspirational District will lead to increased agricultural activity to reduce poverty among tribal communities, making it urgent to understand the relationship between Forest Owlets and agriculture to develop conservation strategies that balance development with the survival of the species.

We investigated the multi-scale habitat selection of Forest Owlets across three nested spatial scales: regional (81 km^2^) encompassing population chunks, landscape mosaics that approximate the extent of their multiple home ranges (4 km^2^), and territory-scale habitat patches (0.25 km^2^). We tested two hypotheses: (H_1_) that Forest Owlet occupancy is higher in tree-based agroforestry than in treeless or intensive agriculture, as this reflects a preference for structurally heterogeneous habitats that provide roosting sites, movement corridors, and accessible prey; and (H_2_) that the forest context modifies the effects of agriculture on occupancy through scale-dependent interactions, where agriculture provides suitable habitat only when it is embedded within landscapes that maintain sufficient forest connectivity. Our objectives were to (1) determine how different types of agricultural land use affect Forest Owlet occupancy, by comparing dry agriculture with trees, without trees, and intensive agriculture systems; and (2) to test whether the forest context modulates the effects of agriculture through interactions between forest and agriculture across spatial scales.

### Study area

The study was conducted in The Dangs District (20°33’–21°05’N, 73°27’–73°56’E), Gujarat, India, which encompasses approximately 1,764 km^2^ of the northern Western Ghats foothills (Fig. 1). The district lies at the westernmost edge of the Forest Owlet’s known range, with the species first confirmed from Gujarat in November 2014 when two individuals were documented in Purna Wildlife Sanctuary (Patel et al., 2015). Elevation ranges from 100 to 1,100 m above sea level, with undulating terrain characterized by rocky lateritic soils.

**FIG. 1.**
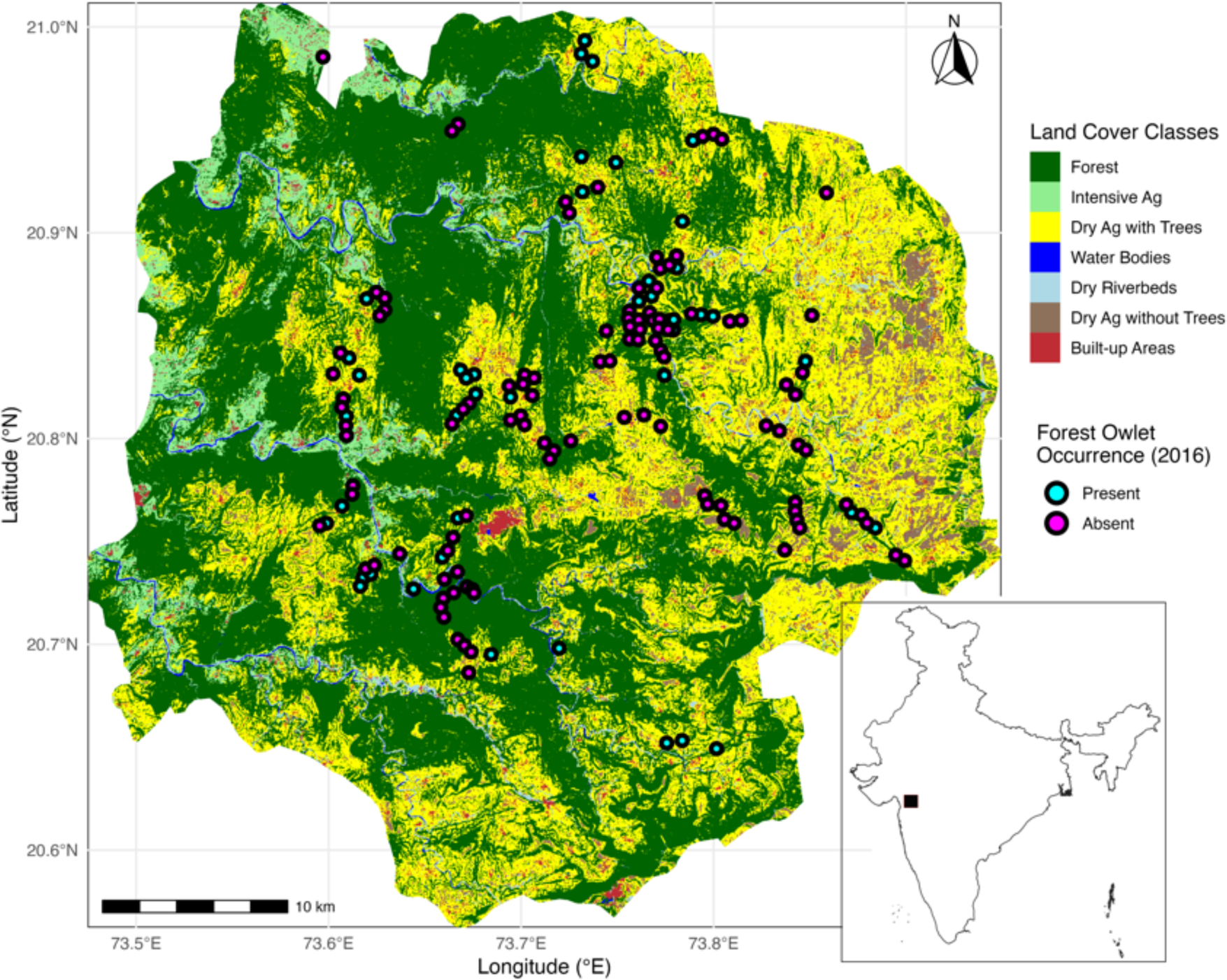
Map of the study area in The Dangs District, Gujarat, India, showing the distribution of surveyed grid cells overlaid on land-cover classification. Inset shows the location of The Dangs District within India and the broader distribution range of the Forest Owlet. The study area encompasses approximately 1,764 km^2^ of the northern Western Ghats foothills.

Monsoon in Dang brings a mean annual rainfall of 2,000–2,500 mm, mostly falling between June and September, the highest in Gujarat. However, the region still experiences seasonal water scarcity due to rocky and hilly terrain. The area is mainly covered by southern tropical dry deciduous forest, which is dominated by teak (*Tectona grandis*), and also features Mahua (*Madhuca indica*), Haldu (*Haldina cordifolia*), and Khair (*Acacia catechu*), Sadad (*Anogeissus latifolia*) other *Terminalia sp* along with bamboos *(Bambusa arundinacea, Dendrocalamus strictus)* as significant species. Forest areas are mixed with agricultural land and human settlements, mainly inhabited by indigenous tribal communities. Human population density in the area is relatively low, at around 129 people per square kilometer, compared to the national average of 382 people per square kilometer (Census of India, 2011).

The Malki agroforestry system is a distinctive feature of the Dangs landscape. Under this state-driven forest-use system, landholders can legally harvest trees on their land, but they must carry out compensatory plantations, which indicates substantial livelihood benefits alongside tree retention (Dasa et al., 2022). This system creates natural variation in agricultural tree density, ranging from traditional agroforestry with dense tree cover to treeless agriculture in drier areas and intensive agriculture where irrigation facilities are available, providing analytical leverage for testing how tree retention affects Forest Owlet occupancy.

## Methods

### Survey design

These surveys were the first conducted after the first confirmed sighting of the species in Gujarat in November 2014 (Patel et al., 2015, 2017). They took place from November 2015 to February 2016, which is the post-monsoon season. During this time, Forest Owlets are more active and vocal, making them easier to detect. Each grid was surveyed using a combination of walking and motorbike transects, covering three to four transects of approximately 4 km each within the 4 km^2^ grid. At intervals of approximately 500 m along each transect, pre-recorded Forest Owlet calls were broadcast for one minute, followed by ten minutes of listening for vocal responses. Both direct sightings and vocal responses were recorded as indicators of presence. Each grid was surveyed three times, at 15-day intervals, to account for temporal variation in detection probability.

### Land cover classification

Land cover was classified using Sentinel-2 imagery from 2016, with cloud masking applied to remove contamination. We computed spectral indices, such as NDVI, NDBI, MNDWI, and BSI, and texture features, including contrast, correlation, energy, and homogeneity. These were incorporated into a composite image, alongside elevation and slope data from the ALOS Digital Elevation Model. Training points were collected for six land-cover classes, with built-up areas initially masked using the Global Human Settlement Layer (Pesaresi, 2023). A Random Forest classifier with 100 trees was trained on the normalized composite image. The final classification achieved an overall accuracy of 81.3% and a Kappa coefficient of 0.776, indicating substantial agreement.

The final land-cover classification consisted of seven classes: (1) Forest, which included dense vegetation areas; (2) Dry agricultural fields with trees, representing traditional Malki agroforestry, with cultivated areas having sparse vegetation and isolated tall trees; (3) Dry agricultural fields without trees, which were cultivated areas with minimal vegetation and no tall trees; (4) Intensive agriculture, characterised by areas with year-round farming activities supported by irrigation, showing a bimodal annual NDVI trend; (5) Water bodies, comprising rivers and lakes; (6) Dry riverbeds, which were areas with exposed rocks and dry river channels; and (7) Built-up areas, including urban and village areas, which were reintroduced using the Global Human Settlement Layer.

### Multi-scale occupancy modelling

We implemented a spatially hierarchical occupancy model following (Lipsey et al., 2017)) to examine Forest Owlet habitat selection across three nested spatial scales (Fig. 2). The broad scale (81 km^2^) captured regional composition and suitability. The intermediate scale (4 km^2^) approximated multiple home-range extents and captured landscape-level forest-agriculture mosaics. The fine scale (0.25 km^2^) captured local habitat patch structure including scattered trees within agricultural fields. The sampling structure comprised 17 broad-scale regions, 71 intermediate-scale landscapes nested within those regions, and 199 surveyed fine-scale sites nested within landscapes, totalling 6,219 spatial units across the three hierarchical scales.

**FIG. 2.**
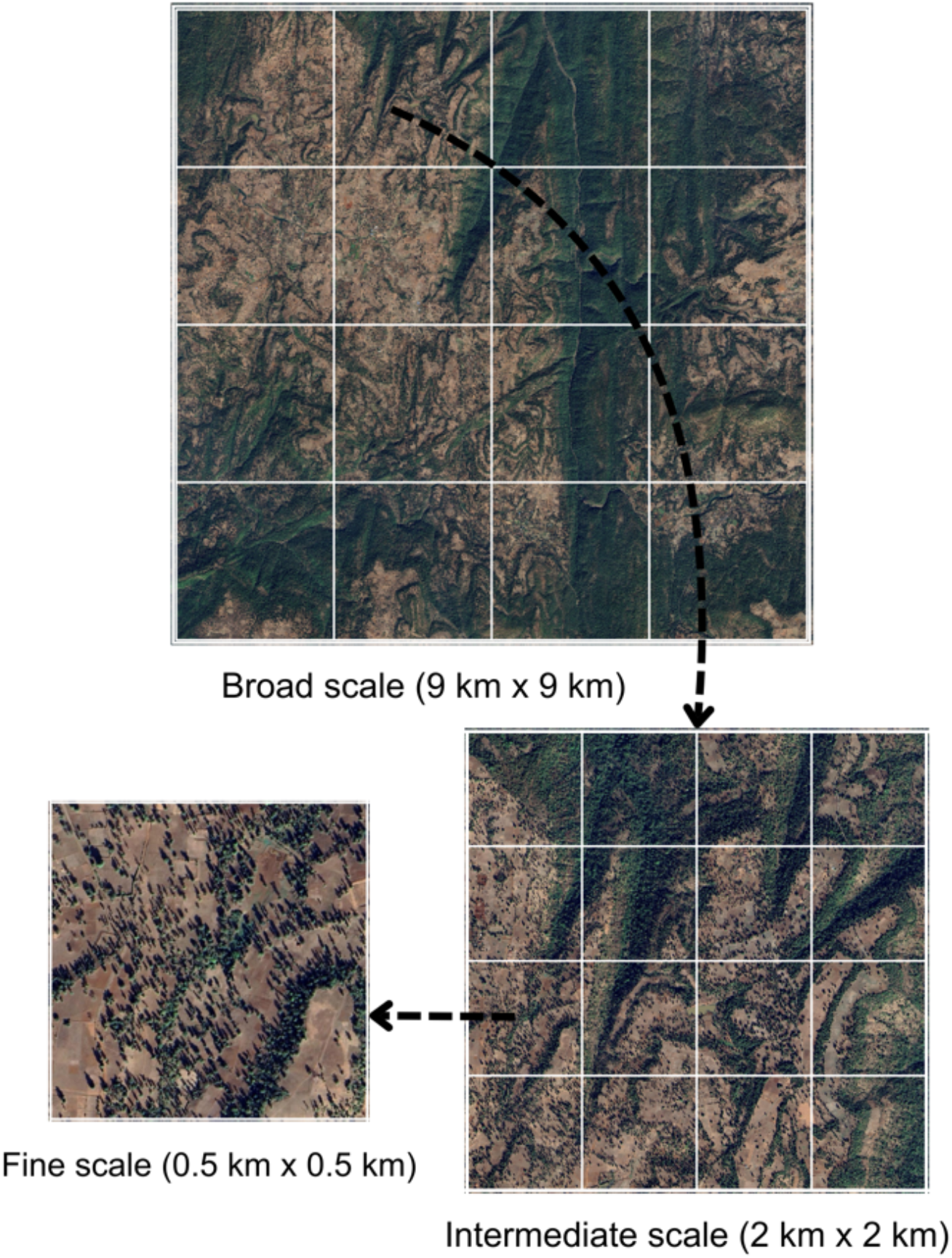
Conceptual framework for the spatially hierarchical occupancy model. The three-scale nested sampling design captures regional landscape composition (81 km^2^, n = 17 broad-scale units), landscape-level forest-agriculture mosaics (4 km^2^, n = 71 intermediate landscapes), and local habitat patch structure (0.25 km^2^, n = 199 fine-scale sites). Yellow arrows illustrate the conditional hierarchical structure where overall occupancy probability is the product of scale-specific occupancies: Ψ = ψ × θ × φ.

The hierarchical model structure allows the overall occupancy probability to be expressed as the product of scale-specific occupancies: Ψ = ψ × θ × φ, where ψ represents the broad-scale occupancy (the probability that a regional unit is occupied), θ represents the intermediate-scale occupancy conditional on broad-scale presence (the probability that a landscape is occupied given that the region is occupied), and φ represents the fine-scale occupancy conditional on intermediate-scale presence (the probability that a site is occupied given that the landscape is occupied). This conditional structure explicitly models hierarchical habitat filtering, where regional-scale selection constrains the landscape-level options, which in turn determines the local site use.

We fitted two model types. The first model focused on the main effects of three agriculture types: dry agriculture with trees, dry agriculture without trees, and intensive agriculture. It did not include interactions. The second model tested the interactions between forest and dry agriculture with trees at each scale, using the proportional cover of each habitat type and their interaction as predictors. All parameters were assigned uninformative uniform priors: β ∼ Uniform(-10, 10). Models were fitted using Bayesian inference in JAGS (Plummer, 2003) via the R2jags package (Su & Yajima, 2024). We ran three Markov chain Monte Carlo chains for 100,000 iterations with a 2,000-iteration burn-in period. Convergence was assessed using the Gelman-Rubin diagnostic 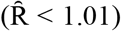, and model fit was compared using the Deviance Information Criterion (Gelman & Hill, 2006).

## Results

### Hierarchical occupancy patterns

Forest Owlets showed a hierarchical pattern of habitat selection across three spatial scales (Table 1). At the regional scale (81 km^2^), the average occupancy of broad units was 67%(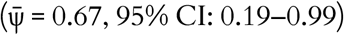, indicating they were widespread but unevenly distributed across the study area. Within the regions that were occupied, the occupancy at the landscape level was 50% 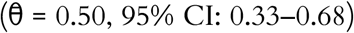, showing that the birds selectively used landscapes based on their habitat con3guration. At the site level, within occupied landscapes, the occupancy was high 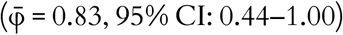, indicating that most suitable patches were used once the landscape context was favourable. This hierarchical 3ltering resulted in an overall occupancy rate of about 28% across the surveyed sites (0.67 × 0.50 × 0.83 ≈ 0.28). The probability of detecting the owlets was moderate 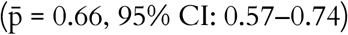.

**TABLE 1.**
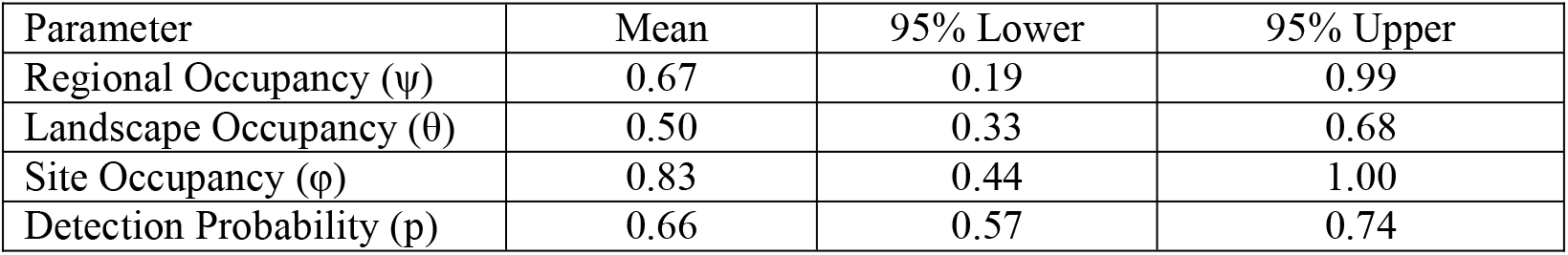
Parameter estimates for the hierarchical multi-scale occupancy model for Forest Owlet. Estimates include the posterior median and 95% Credible Intervals for occupancy at the regional scale (ψ), landscape scale (θ), and site scale (φ), as well as detection probability (p).

The broad credible interval at the regional scale reflects the substantial heterogeneity of landscapes, with some regions having extensive forest-agriculture mosaics that support high occupancy, while others, dominated by intensive agriculture or degraded forest, show low occupancy. These patterns demonstrate top-down filtering, where the composition of regional habitats constrains landscape occupancy, which in turn determines how local sites are used.

### Scale-dependent forest-agriculture interactions

The interaction between forest cover and dry agriculture with trees showed striking scale-dependent patterns that reversed direction across spatial scales (Table 2, Fig. 3). At the broad scale (81 km^2^), a significant negative interaction was detected (β = -6.82, 95% CI: -9.87 to -1.59), indicating antagonistic effects where regions dominated by agroforestry showed higher occupancy than landscapes dominated by forest. The main effects of forest (β = 4.87, 95% CI: -4.47–9.78) and dry agriculture with trees (β = 3.46, 95% CI: -4.48– 9.26) were not significant on their own, but the negative interaction showed that landscapes with high proportions of both habitat types had lower occupancy than expected from additive effects alone.

**TABLE 2.**
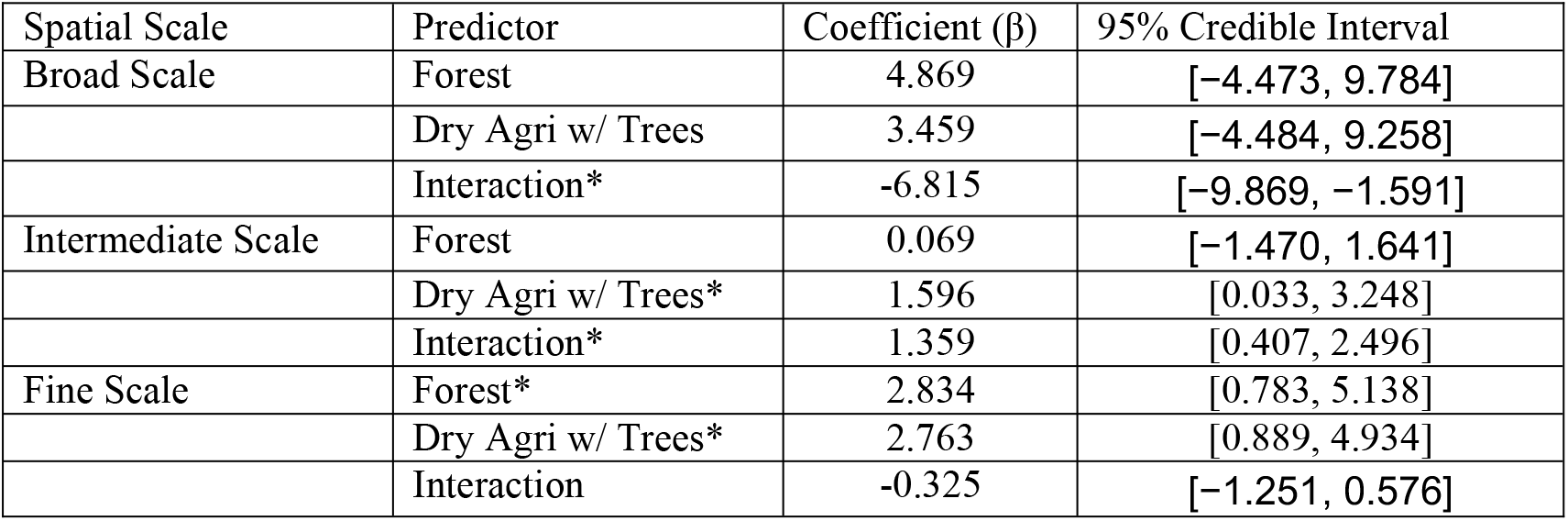
Parameter estimates for the influence of forest and dry agriculture with trees proportions on species occupancy across three spatial scales. The model included an interactive term for both habitat types. Values are the posterior median coefficients β with 95% Credible Intervals in brackets. * indicates coefficients where the 95% Credible Interval does not include zero.

**FIG. 3.**
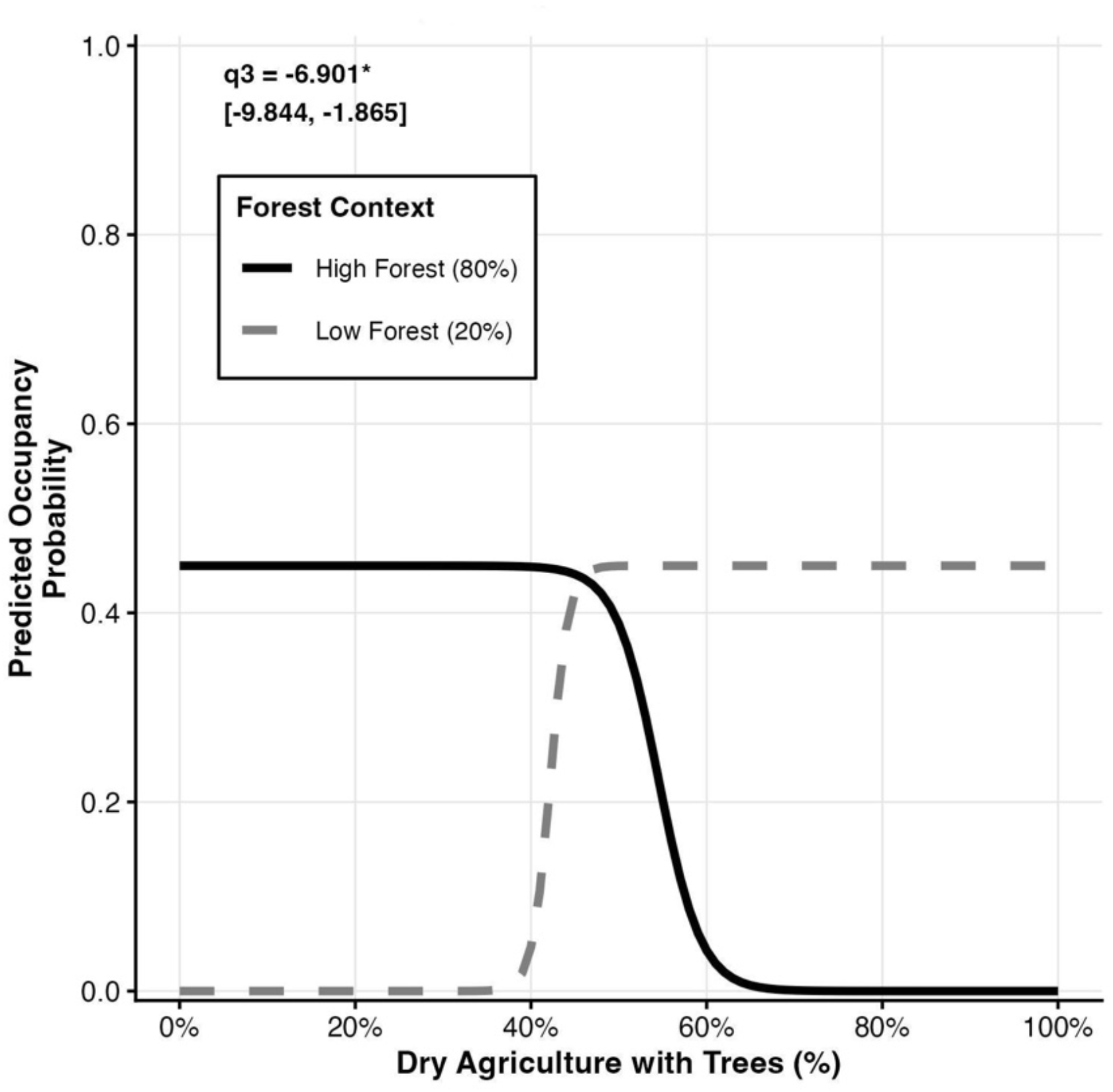
Forest Owlet occupancy probability at the broad scale (81 km^2^) as a function of dry agriculture with trees, stratified by forest cover context. The interaction plot displays predicted occupancy probability (y-axis) against the percentage of dry agriculture with trees (x-axis) for landscapes with high forest cover (80%, solid black line) and low forest cover (20%, dashed grey line). The significant negative interaction (β = -6.901, 95% CI:[-9.844, -1.865]) reveals contrasting effects.

This pattern was reversed at the intermediate scale (4 km^2^), where a significant positive interaction was found (β = 1.36, 95% CI: 0.41–2.50), demonstrating that mosaics of forest and agriculture provided synergistic benefits that exceeded the sum of individual habitat contributions (Fig. 4). The main effect of dry agriculture with trees was also significantly positive (β = 1.60, 95% CI: 0.03–3.25), whereas the main effect of forest was not significant (β = 0.07, 95% CI: -1.47–1.64). At the fine scale (0.25 km^2^), the effects were additive, with no significant interaction (β = -0.33, 95% CI: -1.25–0.58). Both forest (β = 2.83, 95% CI: 0.78–5.14) and dry agriculture with trees (β = 2.76, 95% CI: 0.89–4.93) showed significant positive main effects, indicating that both habitat types provided complementary resources at the territory scale.

**FIG. 4.**
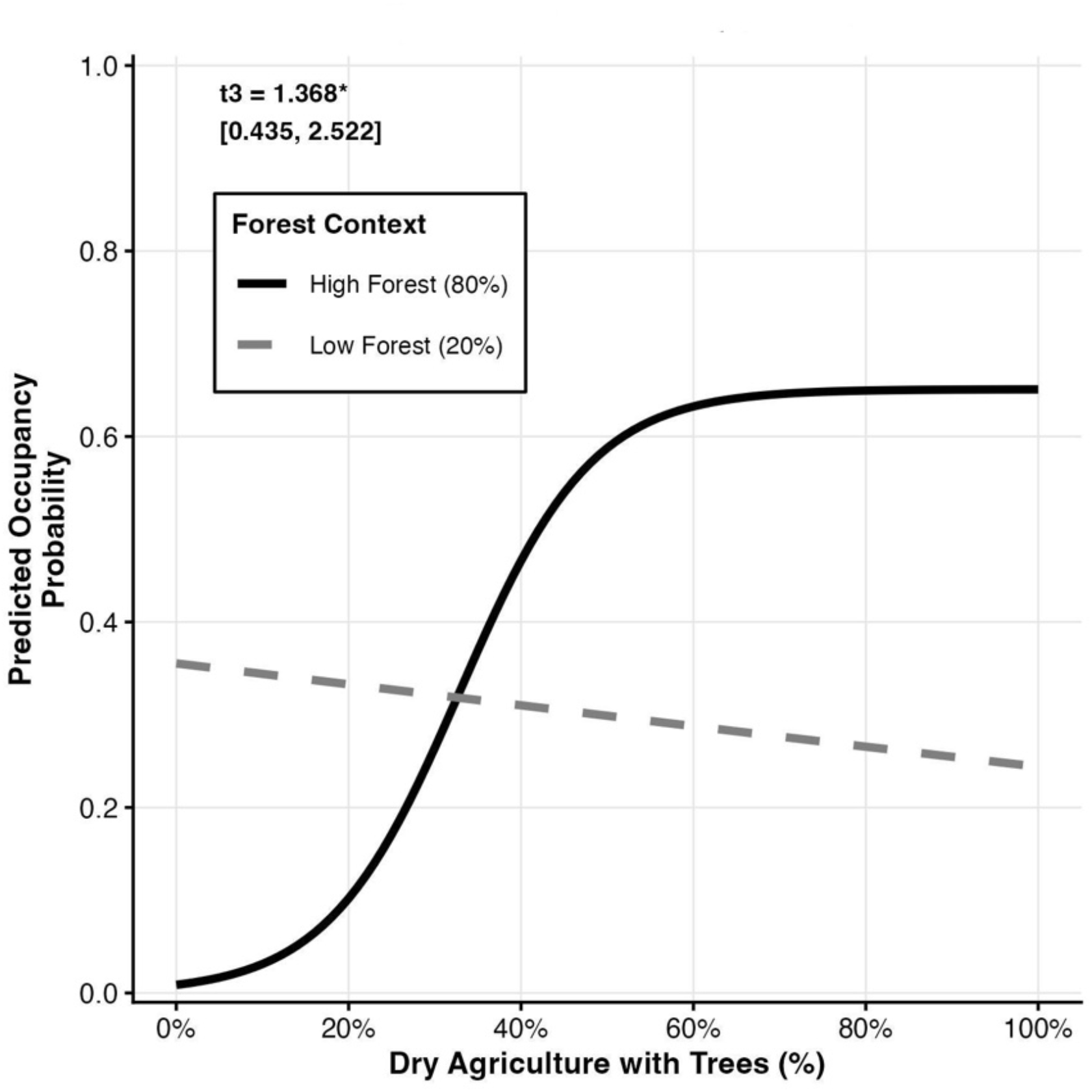
Forest Owlet conditional occupancy probability at the intermediate scale (4 km^2^) as a function of dry agriculture with trees, stratified by forest cover context. The plot shows predicted conditional occupancy probability θ (y-axis) within broad-scale occupied regions against the percentage of dry agriculture with trees (x-axis) for high forest cover (80%, solid black line) and low forest cover (20%, dashed grey line). The significant positive interaction (β = 1.368, 95% CI: [0.435, 2.522]) demonstrates that forest and agroforestry provide synergistic benefits at the landscape scale. Note: these probabilities are conditional on broad-scale occupancy (ψ) and must be multiplied by ψ to obtain overall occupancy.

### Differential effects of agriculture types

Agriculture type comparisons revealed contrasts in habitat suitability, with effects concentrated at the landscape scale (Table 3, Fig. 5). At the intermediate scale (4 km^2^), dry agriculture with trees had a strong positive effect on conditional occupancy (β = 1.17, 95% CI: 0.43–2.02), whereas dry agriculture without trees had a significant negative effect (β = -1.19, 95% CI: -2.28 to -0.29). Intensive agriculture showed no significant relationship (β = 0.09, 95% CI: -1.11–1.15). These patterns suggest that retaining trees in agricultural landscapes is critical for the Forest Owlet’s persistence at the landscape scale, with agroforestry systems providing essential habitat and treeless agriculture acting as an ecological sink habitat.

**TABLE 3.** Parameter estimates for the main effect of three distinct agriculture types (dry agriculture with trees, dry agriculture without trees, and intensive agriculture) on species occupancy across three spatial scales. Values are the posterior median coefficients β with 95% Credible Intervals in brackets. * indicates coefficients where the 95% Credible Interval does not include zero.

| Spatial Scale | Predictor | Coefficient ($\beta$) | 95% Credible Interval |
| --- | --- | --- | --- |
| Broad Scale | Dry Agri w/ Trees | -2.19 | [-8.518, 5.189] |
|  | Dry Agri w/o Trees | 0.403 | [-3.754, 5.384] |
|  | Intensive Agri | -2.889 | [-9.137, 3.800] |
| Intermediate Scale | Dry Agri w/ Trees* | 1.169 | [0.428, 2.018] |
|  | Dry Agri w/o Trees* | -1.187 | [-2.281, -0.292] |
|  | Intensive Agri | 0.094 | [-1.107, 1.152] |
| Fine Scale | Dry Agri w/ Trees | 0.324 | [-0.378, 1.033] |
|  | Dry Agri w/o Trees | 0.003 | [-1.856, 1.978] |
|  | Intensive Agri | -3.81 | [-8.932, 0.300] |

**FIG. 5.**
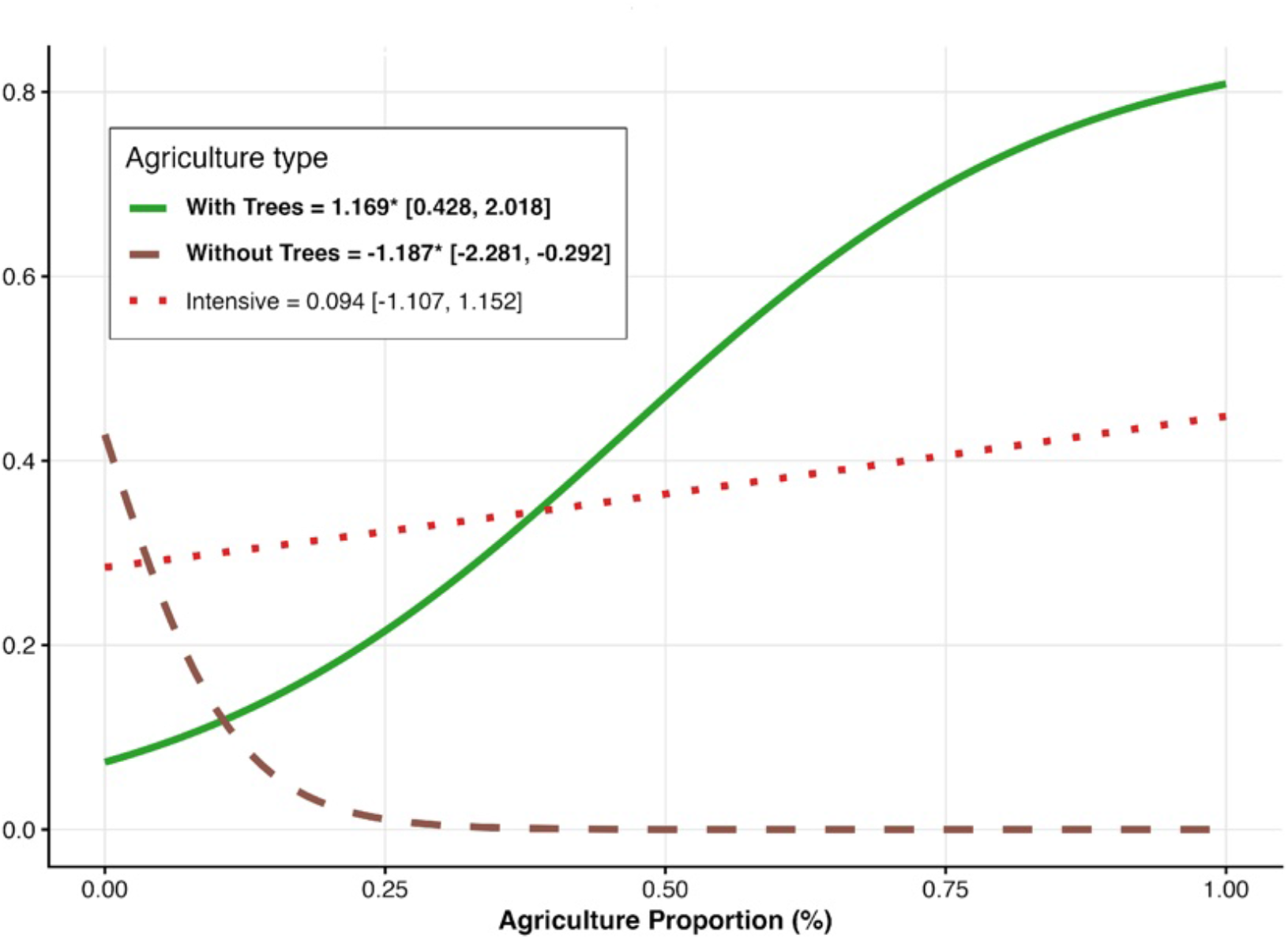
Forest Owlet conditional occupancy probability at the intermediate scale (4 km^2^) in response to different agriculture types. Lines show predicted conditional occupancy (θ) as agriculture proportion increases from 0 to 100%, with all other variables held at mean values. Green solid line = dry agriculture with trees (β = 1.169, 95% CI [0.428, 2.018]); brown dashed line = dry agriculture without trees (β = -1.187, 95% CI [-2.281, -0.292]); red dotted line = intensive agriculture (β = 0.094, 95% CI [-1.107, 1.152]). Asterisks denote coefficients whose 95% credible intervals exclude zero. Note: probabilities are conditional on broad-scale occupancy.

At the broad scale (81 km^2^), none of the agriculture types had significant main effects, although dry agriculture with trees (β = -2.19, 95% CI: -8.52–5.19) and intensive agriculture (β = -2.89, 95% CI: -9.14–3.80) showed a negative trend, while dry agriculture without trees was neutral (β = 0.40, 95% CI: -3.75–5.38). At the fine scale (0.25 km^2^), dry agriculture with trees showed a weak positive but non-significant trend (β = 0.32, 95% CI: -0.38–1.03), dry agriculture without trees was neutral (β = 0.003, 95% CI: -1.86–1.98), and intensive agriculture trended strongly negative, although the credible interval included zero (β = -3.81, 95% CI: -8.93–0.30).

## Discussion

Our hierarchical occupancy analysis showed that Forest Owlets have complex, scale-dependent habitat selection, which brings more clarity to our existing understanding of this endangered species’ habitat requirements. Rather than being specialists of the forest or solely dependent on agricultural fields, Forest Owlets occupy landscapes that are dominated by agroforestry, where agriculture rich in trees and forest patches combine to form heterogeneous mosaics. The reversal of forest-agriculture interaction effects from negative at regional scales to positive at landscape scales shows that habitat suitability cannot be characterised at a single spatial scale, a finding with significant implications for conservation planning in human-modified landscapes.

The significant negative interaction between forest and agroforestry at the broad scale (81 km^2^) indicates that Forest Owlets favour areas where one habitat type is predominant, rather than where both occur in equal proportions. This regional filtering probably shows that the species prefer intact habitats, like forests or agroforestry, instead of mixed fragments. However, within suitable areas at a regional level, the significant positive interaction at the landscape scale (4 km^2^) reveals that habitat complementation becomes essential: forest patches provide cavities for roosting and nesting, while adjacent agroforestry offers accessible foraging opportunities in fallow fields with scattered trees (Ishtiaq & Rahmani, 2005; Yosef et al., 2010; Jathar & Rahmani, 2012).

This cross-scale interaction reversal has been documented in other owl species, but rarely with explicit hierarchical modelling. (Martínez & Zuberogoitia, 2004)) found that Long-eared Owls in Spain avoided extensive continuous forest at landscape scales, while they selected forest patches at home-range scales, a pattern strikingly similar to our Forest Owlet results. Similarly, (Comfort et al., 2016)) showed that Northern Spotted Owls had scale-dependent edge responses, being resilient to hard edges at broad scales but avoiding them at foraging scales. Our findings extend this body of evidence to an endangered Indian endemic, demonstrating that scale-dependent habitat selection may be a widespread phenomenon among owl species that occupy forest-agriculture interfaces.

The strong differential effects of different agriculture types at the landscape scale have direct relevance to conservation. The significant positive effect of dry agriculture with trees (β = 1.17) was in sharp contrast to the significant negative effect of dry agriculture without trees (β = -1.19), showing that the ecological value of agricultural land depends critically on whether trees are retained. This finding supports our first hypothesis that Forest Owlet occupancy is higher in tree-based agroforestry than in treeless or intensive agriculture. The pattern is consistent with the observations of (Patel et al., 2017), who found that 71.4% of Forest Owlet detections were in teak-dominated agricultural landscapes, and with (Khan et al., 2023), who documented positive associations with both high tree density and agricultural cover. The neutral effect of intensive agriculture (β = 0.09) suggests that irrigated year-round cultivation does not strongly attract or repel Forest Owlets at landscape scales, although the negative trend at a finer scale (β = -3.81) indicates that Forest Owlets tend to avoid intensive agriculture at the territory level.

Our results suggest a connection between Forest Owlet ecology and the Malki agroforestry system. The Malki practice maintains a tree-rich agricultural matrix, which Forest Owlets need at landscape scales, by incentivising tree retention and compensatory plantations. Between 1994 and 2019, the Malki system led to a 74% increase in green cover trends in the Dangs forest area, with around one million trees being planted (Dasa et al., 2022). However, we propose the idea of ‘conditional compatibility’ rather than assuming an automatic synergy between the Malki system and Forest Owlet conservation. Although the Malki system helps maintain green cover at a coarse scale, it does not explicitly protect the specific habitat features that are critical for Forest Owlets, such as mature cavity-bearing trees for nesting and roosting.

The hierarchical occupancy structure we documented (regional ψ = 0.67, landscape θ = 0.50, site φ = 0.83) has important implications for spatial conservation prioritisation. The expected occupancy in areas that meet all three scale-specific requirements is approximately 28%, but this figure hides substantial variation. In areas that are unsuitable at a regional level (low ψ), the expected occupancy drops to less than 2%, regardless of the local habitat quality, which is a pattern that analyses at a single scale would miss. Conservation efforts should therefore focus on landscapes where all three scales align, as the expected occupancy here is 5–10 times higher than in areas that are unsuitable at a regional level. Within these priority regions, maintaining mosaics of forest and agriculture at a landscape scale and protecting mature trees in agroforestry fields would address the scales at which habitat selection is strongest.

Several caveats warrant consideration. First, our limited sample size at the broad scale (n = 17 regional units) results in wide credible intervals for regional occupancy (0.19–0.99), potentially limiting what we can infer about regional patterns. Also, our inference of the negative effect of intensive agriculture has wide credible intervals, suggesting a requirement for more focused study on how intensive and mechanised agriculture affects the forest owlet population. Second, the fact that our detection method relied on vocal responses to playback means it may underestimate true occupancy if some individuals do not respond. However, our moderate detection probability (p = 0.66) and three repeat surveys per site provide reasonable confidence in our occupancy estimates. Third, the hierarchical model assumes that occupancy at finer scales is conditional on presence at broader scales, which is biologically reasonable for a territorial species.

Our findings have direct management implications for Forest Owlet conservation in The Dangs and potentially across the species’ range. Conservation planning should use multi-scale frameworks. These frameworks consider the hierarchical nature of habitat selection. This is better than just focusing on single-scale habitat metrics. Next, we can enhance the Malki agroforestry system for Forest Owlet conservation. This can be done by protecting cavity-bearing trees and small forest patches in Malki holdings. Agricultural intensification poses a major threat at a fine scale. This is especially true when farming with trees changes to farming without trees. We should discourage this in other key Forest Owlet areas by using targeted incentives where Malki practice is not followed. Lastly, regional conservation should focus on keeping the forest-agriculture interface. This can be done by using clear zoning. It helps avoid total afforestation, which removes open foraging areas, and total agricultural conversion, which removes nesting sites.

As The Dangs undergoes agricultural development under its Aspirational District designation, our results provide science-based guidance for sustainable intensification that accommodates Forest Owlet conservation. The species prefers traditional agroforestry landscapes. This offers hope that conservation and development can work together, as long as key habitat elements are kept. Multi-scale habitat assessments approach needs to be applied in other regions of the Forest Owlet’s range in Maharashtra and Madhya Pradesh.

Doing so will help test these scale-dependent patterns and guide conservation planning for this critically endangered species.

## Author contributions

JP and AV conceptualised the idea and designed the study; JP, KG and SP did the data collection and field coordinations; JP analysed the data and prepared the initial draft and KS and AV provided critical feedback and enriched the manuscript through many discussions.

## Acknowledgements

We thank Dhaval Patel and the Vidyanagar Nature Club for project management and logistical support. We are grateful to Mehul, Neeraj, Vishal, and Brinky for volunteering at different stages of the project, and to Dr. Ankit Rathod and Nitesh Gamit for field support. We sincerely thank local communities for sharing their knowledge and honest perceptions during surveys, which transformed our outlook on Forest Owlet conservation from pessimistic to optimistic.

We acknowledge Dr. Dheeraj Mittal (then DCF, South Dangs), Shri Anand Kumar (then DCF, North Dangs), and Shri S. C. Pant (then PCCF & HoFF) for providing necessary permissions. This research was supported by the Rufford Foundation and the Indian Bird Conservation Network (IBCN).

## Ethical standard

Authors have acquired mandatory permission and clearance from the state forest department to carry out this study.

## Data availability

The dataset supporting this study is provided as a CSV file in the supplementary material. To protect the Forest Owlet population from potential disturbance, the precise geographic coordinates (latitude and longitude) have been withheld. The dataset still includes the nested hierarchical grid ID structure and all habitat covariates at broad, intermediate, and fine spatial scales, as well as species detection histories. Any additional data requests should be directed to the corresponding author.

